# Deep Learning provides exceptional accuracy to ECoG-based Functional Language Mapping for epilepsy surgery

**DOI:** 10.1101/497644

**Authors:** Harish RaviPrakash, Milena Korostenskaja, Eduardo M. Castillo, Ki H. Lee, Christine M. Salinas, James Baumgartner, Syed M. Anwar, Concetto Spampinato, Ulas Bagci

## Abstract

The success of surgical resection in epilepsy patients depends on preserving functionally critical brain regions, while removing pathological tissues. Being the gold standard, electro-cortical stimulation mapping (ESM) helps surgeons in localizing the function of eloquent cortex through electrical stimulation of electrodes placed directly on the cortical brain surface. Due to the potential hazards of ESM, including increased risk of provoked seizures, electrocorticography based functional mapping (ECOG-FM) was introduced as a safer alternative approach. However, ECoG-FM has a low success rate when compared to the ESM. In this study, we address this critical limitation by developing a new algorithm based on deep learning for ECoG-FM and thereby we achieve an accuracy comparable to ESM in identifying eloquent language cortex. In our experiments, with 11 epilepsy patients who underwent presurgical evaluation (through deep learning-based signal analysis on 637 electrodes), our proposed algorithm made an exceptional 23% improvement with respect to the conventional ECoG-FM analysis (∼60%). We obtained the state-of-the-art accuracy of 83.05% in identifying language regions, which has never been achieved before. Our findings have demonstrated, for the first time, that deep learning powered ECoG-FM can serve as a stand-alone modality and avoid likely hazards of the ESM in epilepsy surgery. Hence, reducing the potential for developing post-surgical morbidity in the language function.

## 1. Introduction

Epilepsy is a chronic neurological disorder characterized by recurrent, unpredictable seizures with over 65 million reported cases around the world (1). Approximately 20% of these patients are diagnosed with *drug-resistant epilepsy (DRE)*. The only possible treatment in a majority of these cases is surgical intervention. During epilepsy surgery the pathological brain tissue, which is associated with seizures, might be surgically removed. While epilepsy surgery is a curative option for drug-resistant epilepsy, neurosurgeons need to avoid removing tissues associated with language, sensory, and motor functions. This calls for an accurate identification and localization of these functionally significant brain regions. The surgical procedure can be performed more accurately with a precise localization for preventing any corresponding post-surgical neurological/functional deficits. This would ensure an improved and sustainable post-surgical quality of life for patients presented for brain surgery.

Electro-cortical stimulation mapping (ESM) has been considered as the gold standard for functional cortex localization in epilepsy surgery. ESM is an invasive procedure that uses electrodes placed on the surface of the brain (grid electrodes) or within the brain (depth electrodes). It is considered vital for reducing the risk of language deficits post-surgery and therefore, expanding surgical options. ESM has a long history of serving as the main modality for pre-surgical functional mapping of epilepsy patients. Acute electrical cortical stimulation was successfully performed in 1950 during epilepsy surgery by Penfield and colleagues (2,3). During ESM, pairs of electrodes covering the region of interest (in our case - eloquent cortex) are stimulated by delivering a brief electric pulse. The stimulation temporarily disables/inhibits the cortical area of interest (creates a temporary functional lesion). Behavioral changes such as unusual sensation, involuntary movements, or language impairments (i.e., speech paucity), observed during stimulation indicates that the tested area is essential to that task performance, and its resection might lead to functional deficits. Ojemann et al. (4) studied language localization using ESM, on a large dataset of 117 patients. The study found that there was sufficiently large individual variability in the exact location of language function and concluded that there was a need for an improved language localization model. Much later, more standard and effective tasks for expressive language localization, such as verb generation (5) and picture naming (6), were tested with the increased use of ESM.

However, one major drawback of ESM is its potential to induce after-discharges (7), which could result in seizures. Since stimulation provoked seizures can occur rather frequently during ESM procedures (8), ESM tests often need to be repeated, leading to extended time and effort from medical professionals (neuropsychologists and/or neurologists). In some cases, the ESM procedure cannot be completed due to repeated seizure activity and/or its consequences.

The current limitations of the ESM have created a strong need for establishing other independent functional mapping modalities to identify eloquent cortex. Unfortunately, as of now, none of the existing neuro-imaging modalities are flexible enough to provide functional mapping results in real time in the operating room. Therefore, the search for a stand-alone methodology for functional eloquent cortex localization has been continuing and resulted in attempts to use electrocorticography (ECoG) as a viable alternative. ECoG is the invasive version of electroencephalography (EEG) and sometimes also referred to as intracranial encephalography, demonstrating excellent temporal resolution like EEG. Importantly, ECoG equipment is portable and can be utilized both at the patients’ bed-side and intra-operatively. Unlike EEG, it overcomes the problem of poor spatial resolution, since the activity of interest is recorded directly from the cortical brain surface. It also avoids the problem of electrical signal attenuation in EEG caused by the signal propagation through tissues surrounding the brain. To record ECoG signals, a craniotomy (removal of the skull section: *bone flap*) is performed and the dura is opened to access the brain tissue. The arrays of grid of electrodes (**Figure 1a, left**) are then placed on the exposed cerebral cortex. Following this, ECoG-based functional mapping (ECoG-FM) is performed, while task-based responses from grid electrodes are recorded. Since there is no external electrical stimulation during this process, ECoG-FM is considered as a safer alternative to ESM. When performed in real time (9), ECoG-FM procedure can be referred to as real-time functional mapping (RTFM) (10–13). **Figure 1a** demonstrates the general setup for ECoG-FM recordings.

**Figure 1:**
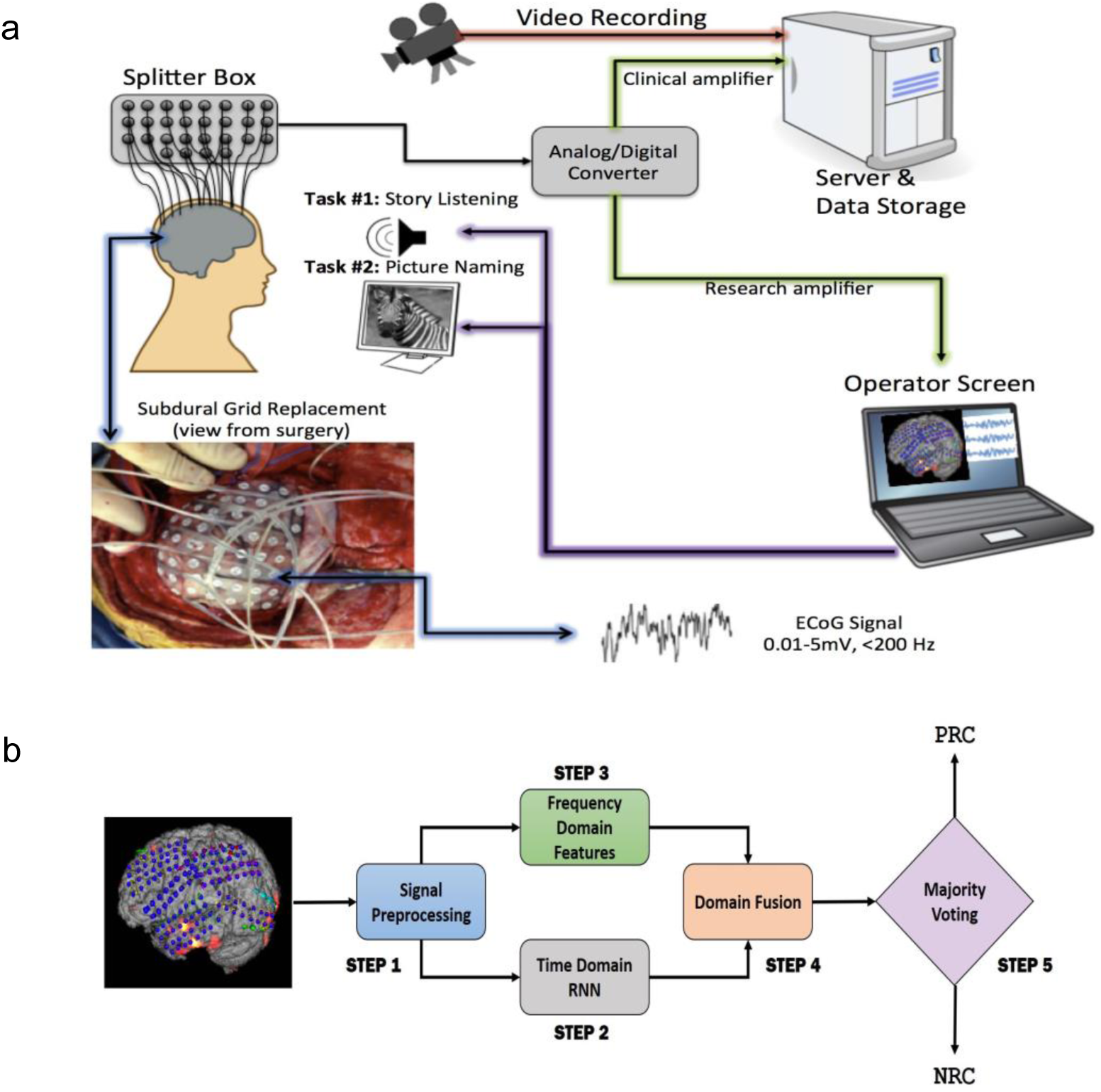
Overview of a) the language localization framework with ECoG-based functional mapping (ECoG-FM) approach. ECoG signal recording, data transfer, storage, research and clinical paths, and tasks are illustrated. ECoG signals are obtained in response to task-related changes (e.g., picture naming) from grid electrodes implanted on the cortical surface in the subdural space, b) the proposed ECoG signal classification approach for each channel on the cortical surface in the subdural space. PRC - Positive Response Channel, NRC - Negative Response Channel. Each individual step is considered as a “module” in the overall system design.

### 1.1 Related works and existing critical challenges

ECoG-based approaches have been used successfully for motor cortex localization (10,11,14–16). In comparison, the localization of functional language cortex appears far more complex and challenging (17). Current localization approaches are based on detecting positive response channels (called *active channels or active electrodes*) among the set of all channels. A baseline recording of each channel at resting-state is used to determine signal characteristics at specific frequency ranges. Most often, power of the ECoG signal lies within the *α, β*, and (primarily) high-*γ* (70Hz-170Hz) frequency bands (9,18). These values are compared with the signal power measured during the execution of language task. The results of this approach for language mapping have not achieved desirable accuracy. For example, Arya et al. (19) studied high-*γ* response from ECoG recording of 7 patients during spontaneous conversation. The results showed low specificity and accuracy. In a follow up study, Arya et al. (20) demonstrated high-gamma modulation for the story-listening task and achieved high specificity, but sensitivity remained low. Korostenskaja et al. (21) showed that similar to the results for motor cortex, ECoG-FM can be used for eloquent language cortex localization as a complimentary technique with ESM, but not as a stand-alone modality. It has also been demonstrated that ECoG-FM can be used as a guiding tool for ESM, thereby reducing the time of ESM procedure and decreasing the risk of provoked seizures (13,22).

Despite their potential, the current ECoG-FM approaches are not found capable enough to be used as a stand-alone methodology for accurate language mapping. To address these challenges and provide ECoG-FM more independence in eloquent language cortex localization, we fill in the following currently existing methodological gaps:

i. Available approaches compare a channel’s signal with its resting-state (baseline) recording and do not compare the channels’ characteristics to other recorded channels.
ii. The signal characteristics in the frequency range beyond high-*γ* band have not been explored yet to the best of our knowledge.
iii. There has been limited work on validation of ECoG-FM for language cortex localization; hence, there are more evidences needed for utilization of ECoG-FM in the clinics.

In our previous works (23,24), we showed the feasibility of utilizing conventional machine learning methods for channel response classification by using the whole signal spectrum (not limited to *α, β* & *high γ*) and without using the baseline recording. This was one of the first machine learning based approaches in this field with strong results and demonstrated the potential of ECoG-FM signal to be analyzed more accurately compared to conventional signal processing based descriptive methods.

### 1.2 Summary of our contributions

We propose an innovative deep learning algorithm for accurately classifying the channel response of eloquent cortex, alleviating the current challenges of ECoG-FM. We also show the effectiveness of using deep learning methods in problems with limited number of patients but sufficient samples to achieve desired performance. Our main contributions are as follows,

1. We have used the complete ECoG signal frequency spectrum for the first time to the best of our knowledge to identify signal characteristics for mapping language cortex.
2. Our innovative deep learning models achieve state-of-the-art performance in language cortex mapping using ECoG-FM. We have achieved 82% sensitivity in classifying both positive and negative response channels.
3. We have shown that deep learning models can be successfully used in studies even with a limited number of subjects and low dimensional (1D) data. The requirement for sufficient data to train our deep learning model is satisfied as we show that the number of data points (due to a dense grid placement and high-resolution signal) are adequate and our experimental results add credence to these findings.
4. Our results have indicated that with the proposed deep learning-based classification model, ECoG-FM can be reliably used as a standalone technique for functional language mapping.

## 2. Materials and Methods

An overview of the proposed system is illustrated in **Figure 1b**. We pre-processed the ECoG signals using temporal filtering such that the selected samples were synchronized with the start and end points of the task resulting in equal length blocks. We then divided the equal length ECoG signal blocks into overlapping sub-blocks of data. Our aim was to learn *discriminative signal patterns* and eventually reduced the computational load (Step 1). We learned different sets of signal features independently: frequency domain (i.e. auto-regression) and time domain (deep learning-based features) in Step 2 and Step 3, respectively. After we combined the learned features, we trained a recurrent neural network (RNN), a class of deep learning algorithm suited for analyzing sequential data, to classify sub-blocks of signals (Step 4). Finally, we used the majority voting technique to combine these sub-block labels and determine an overall (PRC or NRC) channel label (Step 5). In the following sub-sections, we will describe each module of our proposed system in detail.

### 2.1 Dataset

The study was approved by the Institutional Review Board at AdventHealth Orlando, Orlando, USA. We recruited eleven patients with drug-resistant epilepsy, who underwent pre-surgical evaluation with ECoG grid implantation. All patients provided their written informed consent to participate in this study. The patients were teenagers and adults with an average age of 23.18 ± 11.61 years (See Supplementary **Table 1** for summary of the patient demographics).

### 2.2 Pre-processing ECoG Signal (Step 1)

As a first step of preparing the recorded data for deep learning-based analysis, non-task/control time points in the signal were eliminated using temporal filtering. Hence, the spontaneous activity recordings before the start and trailing signals at the end of the experiment were discarded from the blocks. These synchronized and uniform length blocks were fed to the deep networks for functional channel classification (See Supplementary for more details about this step).

### 2.3 Time Domain RNN (Step 2)

Our goal was to find discriminative signal patterns from ECoG signals, which are time varying and non-stationary 1D sequences. They are non-stationary because task based ECoG recordings can have signal statistics which depend on the time relative to the events. Inspired by the effectiveness of recurrent neural networks in sequence classification tasks in different domains (25–27), we have developed RNN based deep neural network algorithms to extract discriminative features from time-domain ECoG data. We hypothesized that limitations of the conventional spectral (frequency-based) or time-based signal analysis methods can be overcome with RNN based methods. In RNNs outputs from previous time steps are taken as input for the current time step, thereby forming a directed cyclic graph. RNNs thus learn the relationships in sequential data thereby retaining higher contextual information.

The ECoG signal consists of over a thousand samples every second; to learn signal characteristics from such a noisy data, we first used popular EEG features (28–30). A sliding window approach was applied to extract features, which were then concatenated into a single feature vector to represent the control and active-task blocks in the signal. The extracted features included mean, skew, kurtosis, peak to peak value, and Hjorth values (details in supplementary).

The time domain features were fed to the learning module illustrated in **Figure 2**. The complete ECoG signal contained both control and active task signals, thus; the sub-blocks of control signals were ignored and the input to this time domain module was sub-blocks of active task signals. Recently, 1D convolutional networks have been shown to perform well in time series forecasting and classification tasks (31,32). We designed the module to have two paths comprising of 1D convolutional layers and long-short term memory blocks. LSTM, introduced in 1997 (33), is a type of RNN that has the ability to learn long-term dependencies of data. In literature, LSTM and its variants have primarily been used in 1D sequence classification tasks (34,35) and prediction tasks (36,37). In our experiments, we used multiple LSTM layers in different exploratory configurations to learn a more complex feature representation of the input signal.

**Figure 2:**
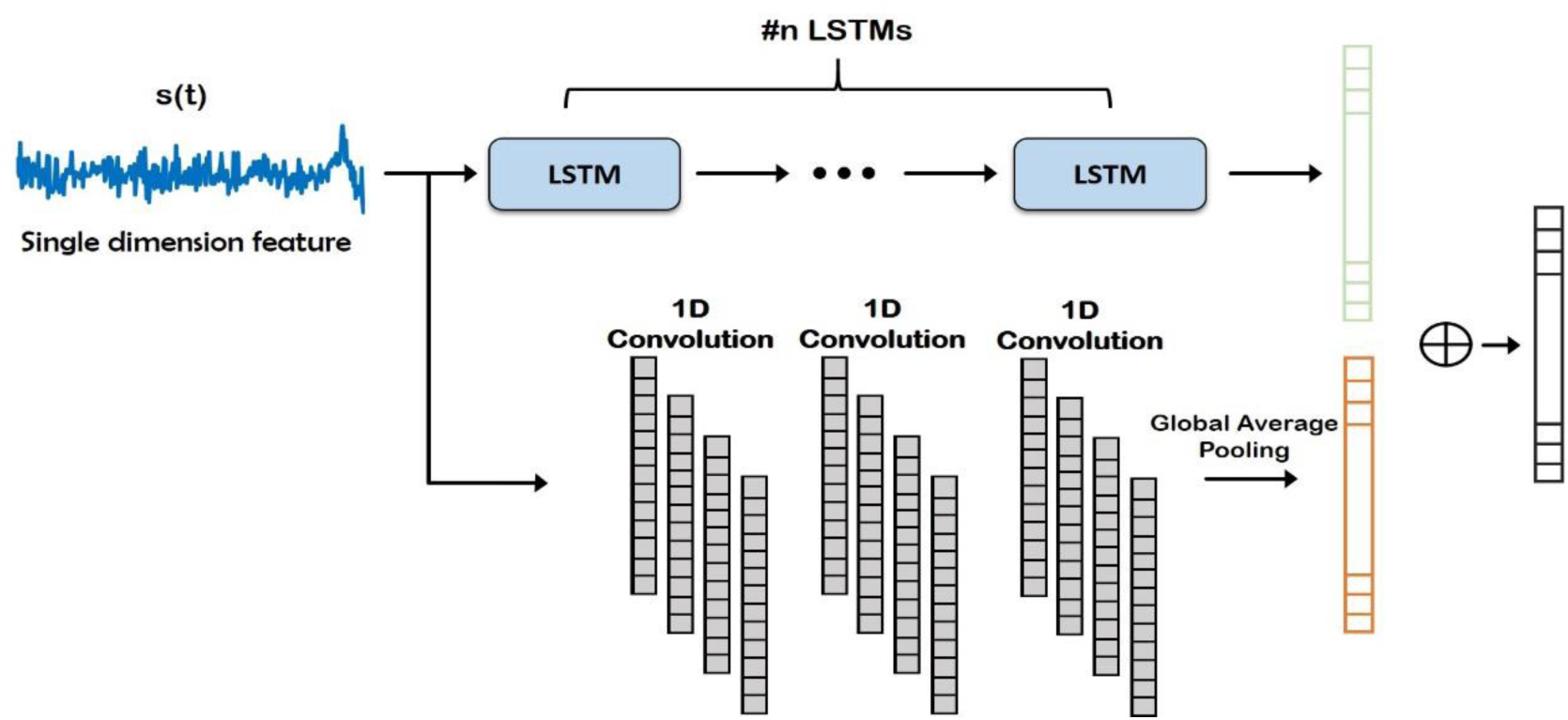
Deep network structure for extraction of time domain features from the input signal. Note that these features are combined with frequency domain features (Auto-Regressive) later for final prediction of the channels.

### 2.4 Frequency Domain Features (Step 3)

One of the objectives of our study was to analyze multi-domain (time and frequency independently) and hybrid-domain (time and frequency combined) signal characteristics. This kind of thorough comparison has never done before to the best of our knowledge for ECoG-FM. In this step (Step 3), we focused on the spectral characterization of signals. Conventional ECoG signal classification approaches are based on frequency-domain, where spectral analysis of the signal is performed to identify the channel response. Traditionally, spectral estimation of the signals is performed by fitting a parametric time domain model to the ECoG signals. One of the most commonly employed models/approaches in this category is the autoregressive model. An AR model for a discrete signal *x*[*n*] is represented as,

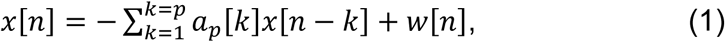

where *a*_*p*_[*k*] are the AR coefficients, *p* is the order of the AR model, *w*[*n*] is a zero mean white noise process with a variance *ρ*. Once the model in Eq. (1) is solved, the resulting AR parameters were used for characterization of the ECoG signal from frequency-domain perspective.

Methods to solve for the AR parameters are diverse, we used the reflection coefficient estimation-based methods (see Supplementary for details of the parameter selection procedure).

### 2.5 Fusion for Hybrid Domain (Step 4)

Using LSTMs in Step 2, we learned a different set of features (i.e., time domain) than the AR features that were generated in Step 3. In domain fusion step (Step 4), these two (largely) complimentary features were combined to obtain a hybrid signal representation model with a new deep network setting, Domain Fusion Network (DFN) (See **Figure 3**). Although the merging of the two feature vectors can be done in multiple ways, we used a concatenation approach to get full benefit of each domain (time vs. frequency).

**Figure 3:**
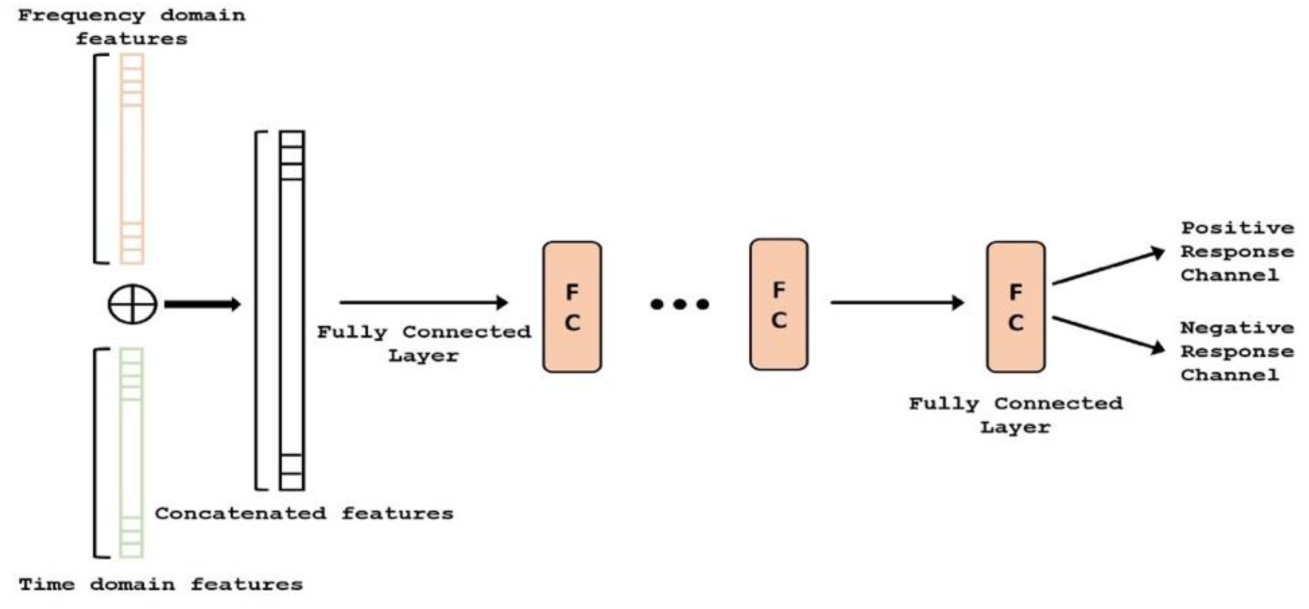
Deep network structure of the fusion module. AR (i.e., frequency) features (orange) and time-domain features (green) are concatenated and classified.

In concatenation, we assumed independence of features; hence, we did not use element-wise multiplication or other approaches for data merging. Since convolution helps identify local patterns and reduce redundant information in the data, the complete feature vector (after concatenation) was then passed through multiple layers of 1D convolutions with an activation function, to weight each feature based on its contribution to the classification problem (PRC vs NRC). Following the 1D convolution layers, the output feature maps were spatially averaged using Global Average Pooling (38), making the DFN more robust to spatial translations of the input data and introducing structural regularization to the feature maps. Finally, we inserted a single fully connected layer into the DFN and used a sigmoid activation to perform the final classification.

### 2.6 Majority Voting (Step 5)

The output of the domain fusion model was a label for the input signal, which was a sub-block. Signal from each channel/electrode was made up of hundreds of sub-blocks of the signals with reasonable overlapping. Therefore, for classifying a channel as either PRC or NRC, we hypothesize that the output that is observed more commonly is assigned as the final label. For this purpose, we apply majority voting on the output for each sub-block. For instance, if a channel included 354 sub-blocks and more than 50% of sub-blocks indicated a positive response, that channel was labelled as a PRC. As a rule, whenever the number of negative and positive responses are equal, the channel will not be assigned any label. Although, we did not observe any such channel in our experiments.

## 3. Experiments and Results

### 3.1 Task Paradigm

ECoG signals from the implanted subdural grids are split into two streams: one for continuous clinical seizure monitoring and the other for ECoG-FM (**Figure 1a**). The tool used to record the incoming ECoG signal was BCI2000(9). A baseline recording of the cortical activity was first acquired to capture the “resting-state” neuronal activity. Following this *baseline* recording step, paradigms similar to those employed in ESM or functional magnetic resonance imaging (fMRI) were used to record the task-related ECoG signal for functional mapping (39). **Figure 4** shows one such paradigm, mimicking the exact details of the experimental setup we have used for the language comprehension task. Alternate 30 second blocks of ECoG data during “control” and “active” conditions were recorded continuously at a fixed sampling rate of 1200 Hz.

**Figure 4:**
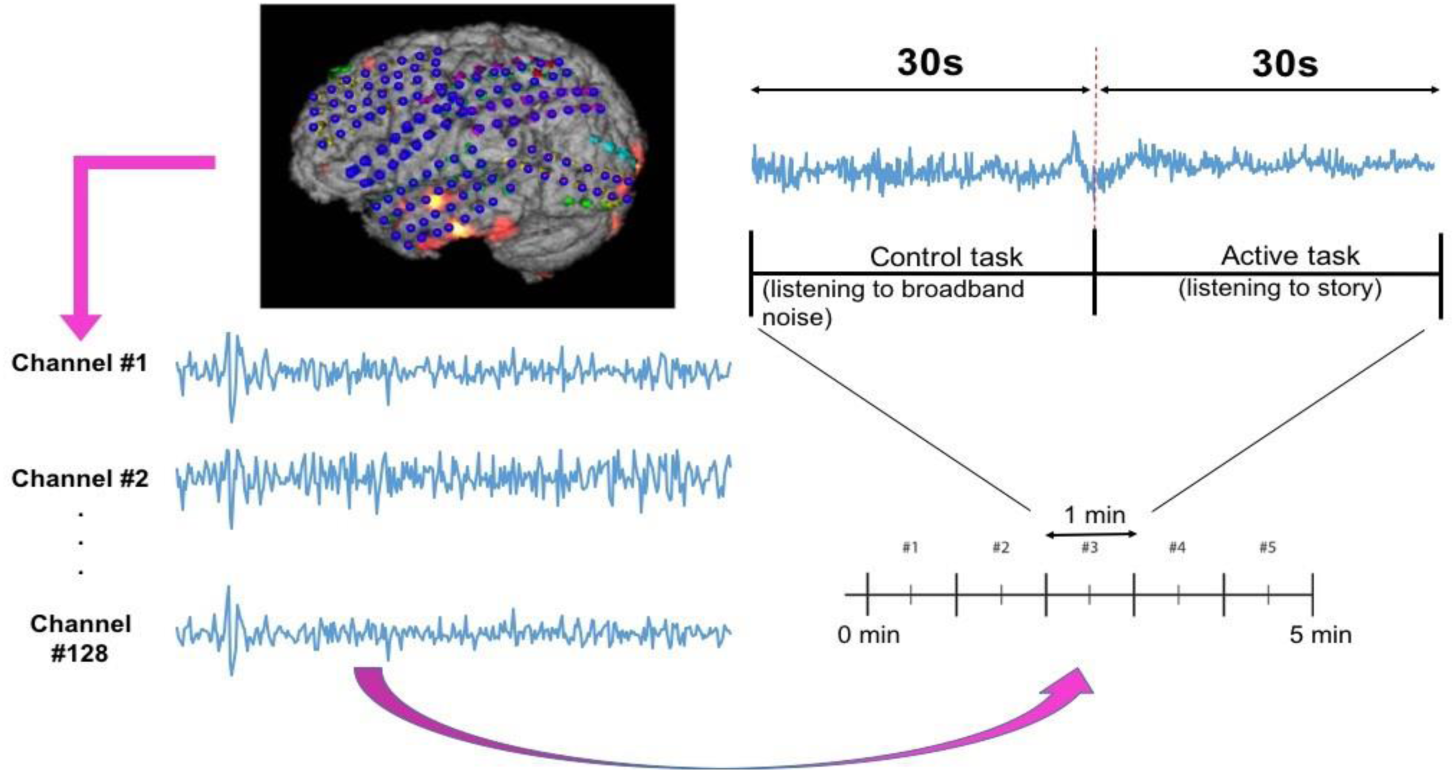
Subdural grid localization and position of ECoG electrodes (128 of them) on the brain surface of individual patient are illustrated (left). For a sample of 1 min language test, signals from both control and active tasks are illustrated (right).

For the language comprehension task, the *active condition* implies listening to a story, while the control task involves listening to broadband noise (40). For the active condition (i.e., listening to a story), a different story was selected for each block in order to keep the patient attentive and responsive. Both control and active sequences would activate sense of hearing, but the story listening task will particularly activate the language function. We hypothesize that this would suffice in eliciting the desired response for mapping eloquent cortex related to language function and our results have verified this hypothesis. For this purpose, the system recorded information from 128 ECoG channels (128 electrodes in **Figure 4**) by using *g*.*USBamp bio-signal amplifiers* (g.tec Medical Engineering GmbH, Austria) with subdural ground and reference electrodes. 9

### 3.2 Training Paradigms

Our overall goal was to successfully (and automatically) identify positive response channels (PRCs) and negative response channels (NRCs) in ECoG-FM data using new machine learning models, specifically based on deep neural networks. The ground truth (i.e., reference standard) was inferred from the gold standard ESM results. Owing to the large imbalance in the number of PRCs and NRCs (NRCs outnumbering PRCs by 3:1), we randomly selected equal number of NRCs to balance the data and avoid potential data imbalance problem when training deep learning models.

Each channel’s signal comprised of blocks of active task data and control data, where each active task block was from a different story. The discriminative power of these stories in the classification task was unknown. There is a possibility that features from one story could play a more significant role than others. Additionally, the discriminative power of any particular feature is unknown. To ascertain the role of these, we divided our experimental evaluation approaches into three main categories for data classification. Each approach depended upon the way active task data was included, and the features used in the training process. This structured experimental procedure helped us in determining the usefulness of each component of the signal and provided insights into the response of brain regions (through channel responses) to different signals. We performed experiments with different features and architectures.

### 3.3 Our Proposed Deep Network Architectures

In task-based experiments, a response is generally expected only during the active task period and not in the control or rest period. We used fully convolutional network (FCN) and long short-term memory (LSTM) architectures in the time domain module, since these have shown success in various time-series classification problems (41,42). We built our network by first analyzing the effect of using time domain features during the active task (represented as Active Time-AT). We tested our proposed time domain module by varying the network. We used an FCN (represented as AT^1^) and then added LSTM module to the network (represented as AT^2^). For frequency domain analysis, we added the auto regressive (AR) features to the frequency domain module by passing it through a fully connected layer (represented as AT-AR^1^). **Figure 5a** shows the architectures (the superscripts indicate the variation within an architecture) including AT^1^, AT^2^ and AT-AR^1^. In the domain fusion module, we tested different combinations of 1D convolutions (represented as AT-AR^2^) and fully connected layers (represented as AT-AR^3^). We also varied the depth of the frequency domain module by adding an additional fully connected layer in the network (represented as AT-AR^4^). The network structures including AT-AR^2^, AT-AR^3^, and AT-AR^4^ are shown in **Figure 5b**. We empirically determined the number of epochs required to train the network such that to avoid overfitting. Our experimental paradigms used time and frequency domain features individually and also in a hybrid manner (combined). We also analyzed the effect of active and control task data.

**Figure 5:**
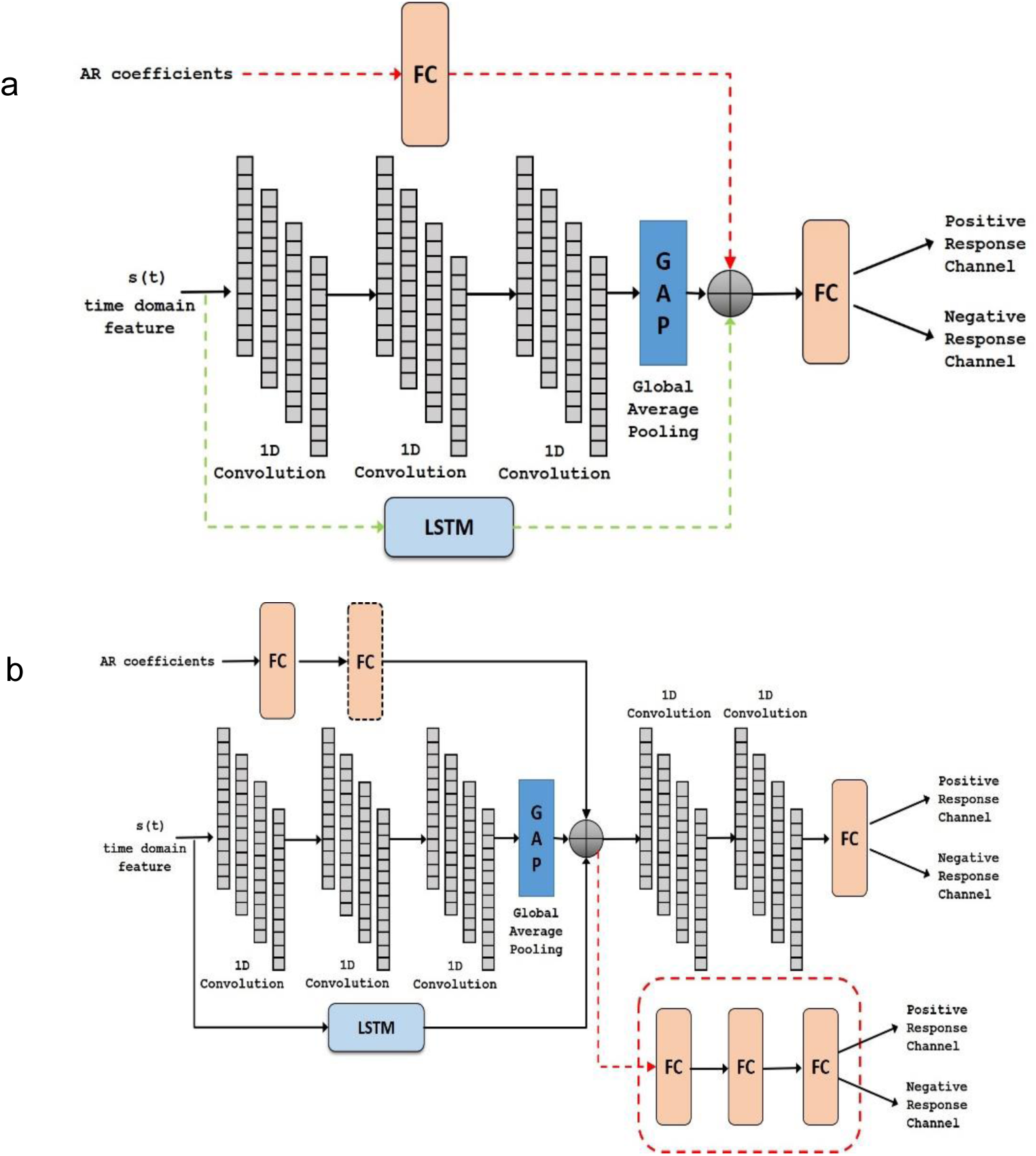
Deep network structure of a) *AT*^1^, *AT*^*2*^ (with added dotted green) and *AT-AR*^*1*^ (with added dotted red and green) and b) *AT-AR*^*2*^, *AT-AR*^*3*^ (replace fusion model with the dotted red model) and *AT-AR*^*4*^ (with added FC in dotted box).

### 3.4 Model Validation

In supervised machine learning approaches, where a model is trained using ground-truth labels, the goal is to maximize predictive accuracy. However, therein lies the risk of memorizing the data rather than learning the optimal features. This problem of memorizing the data or learning the structure of the data to be the noise in the data is often referred to as **overfitting** (43). It is important for a classification model to be able to generalize to unseen data and avoid the problem of overfitting. The method of testing how the analysis/model generalizes to an independent test dataset is known as cross-validation. When a completely independent dataset is not available, as is generally the case, the available data is split into training data and validation/test data. There are different types of cross-validation approaches such as leave-one-out cross-validation, hold-out method, *k*-fold cross-validation, to name a few (44).

### Shuffle-split Cross Validation

Previously, due to the time-consuming nature of training a deep learning model, we applied the hold-out method to validate our proposed models (45). In this method, the model is trained on a part of the available data, while the remaining data is held for testing/validating the model. For effectively testing the generalization and robustness of our proposed models, we validated them using the shuffle-split cross-validation approach. In the shuffle-split cross-validation method, the data is randomly sampled and split into training and testing splits iteratively, similar to the hold-out method. The results are averaged across the number of iterations. This can be seen as repeating the hold-out method k times, such that the data for training and validation is randomly sampled each time. The use of shuffle-split method allows sampling different data combinations rather than a single sampling as in k-fold cross-validation. Since the blocks were randomly assigned to the training and test folds, we ensured that no data from an electrode (channel) was represented in both training and testing folds simultaneously. Hence, if a block of data was assigned to a particular set (training/test), then all blocks belonging to that channel were assigned to the same (training/test) set. This ensured a fair evaluation with better generalization accuracy and helped in avoiding overfitting. For 30-fold cross validation, we repeated the experiments 30 times and used each of these distinct and non-overlapping training-testing sets to evaluate our model accuracies. Prediction accuracy was then calculated by averaging the results of these 30 experiments.

### 3.5 Training on individual features on active task data (Training Paradigm-I)

In this approach, we assumed that the channel response was similar for different stimuli (story) used in this study. Our experimental paradigm consisted of 5 different stories and thus, in this approach, no distinction was made with regards to the story. All of the active task data (i.e., five different story tasks) from a channel were used together for training the network. Among the time domain features (See Section: Methods), we found using random forest method that activity feature gave the best results. Therefore, all our proposed deep learning architecture (**Figure 5**) were first tested using the activity feature (**Table 1**). The addition of LSTM improved the performance of the time domain module. This was further improved by the addition of the frequency domain features using the domain fusion module. We found that increasing the depth of the frequency domain module did not have any obvious benefit in classification performance.

**Table 1:**
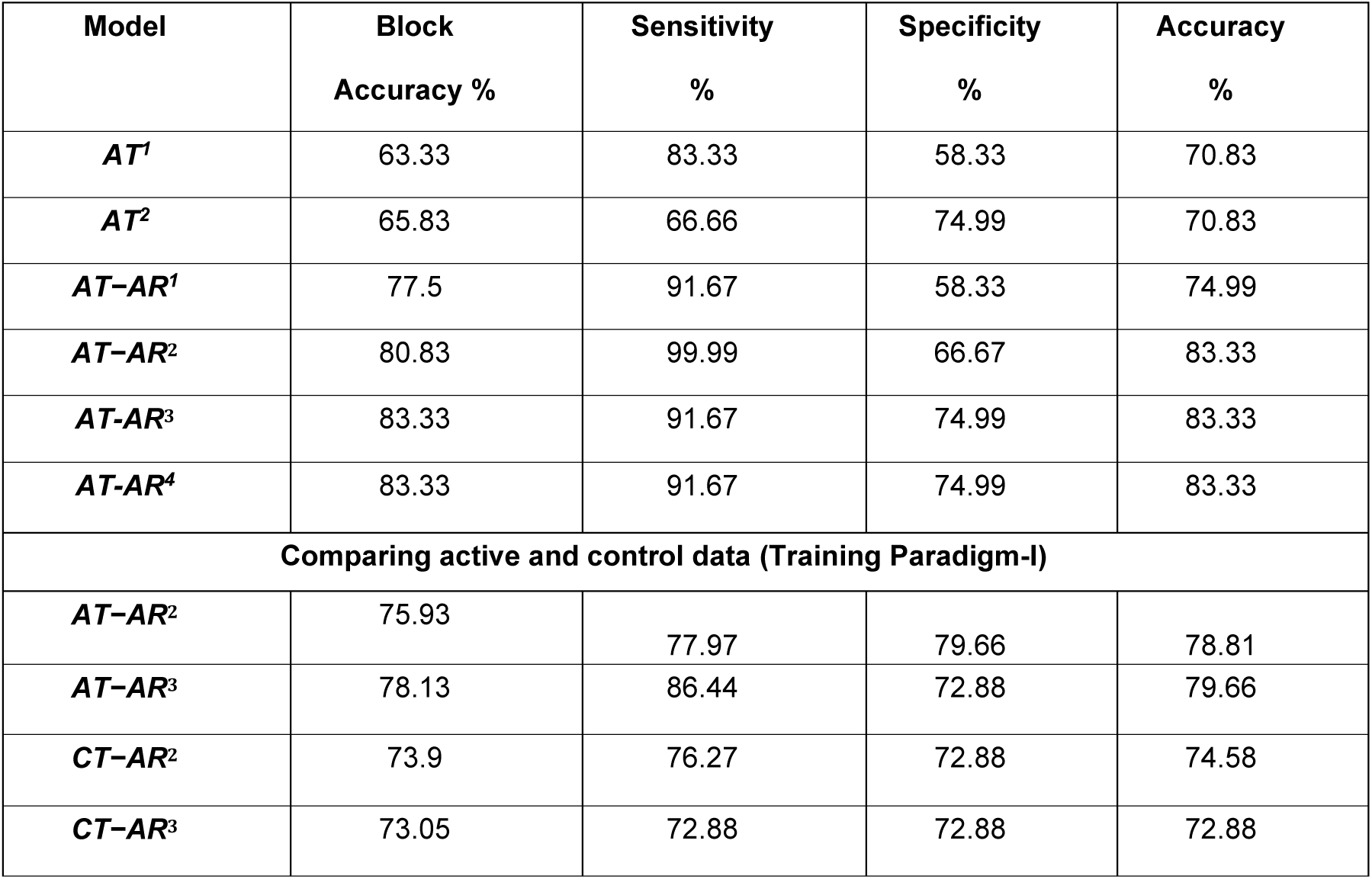
Channel classification accuracy for different network architectures. AT- active time, AR – auto regression, CT- control time.

We also tested our hypothesis that the story listening task (active task) was more discriminative in identifying the eloquent cortex. To compare information present in the active and control task data, we replicated our best performing models (AT-AR^2^ and AT-AR^3^) and fed it with control task data (represented as - CT-AR^2^ and CT-AR^3^, where CT represents control time). We observed that sensitivity of the control data model was lower than that of active data model, indicating a lower discriminative power (**Table 1**) and confirmed our hypothesis. To identify the best features for the channel classification task, we fed the best performing model (AT-AR^3^), with different hand-crafted features and performed cross-validation. The performance with different features was found to be similar (**Table 2**). The mobility feature showed the best performance with high sensitivity and accuracy compared to the other features.

**Table 2:** Channel classification performance parameters (with mean and variance) for active task data with individual hand-crafted time domain features using the AT-AR^3^ model (training paradigm-I).

| Features | Sensitivity %<br>$\mu \pm \sigma$ | Specificity %<br>$\mu \pm \sigma$ | Accuracy %<br>$\mu \pm \sigma$ |
| --- | --- | --- | --- |
| *Mean* | 83.33 ± 10.09 | 81.11 ± 9.61 | 82.22 ± 6.71 |
| *Skew* | 81.67 ± 9.47 | 81.67 ± 9.95 | 81.67 ± 5.54 |
| *Kurtosis* | 82.78 ± 11.97 | 79.17 ± 10.03 | 80.97 ± 7.81 |
| *P2P* | 82.22 ± 10.91 | 80.00 ± 9.77 | 81.11 ± 7.11 |
| *Activity* | 82.22 ± 9.31 | 79.44 ± 10.70 | 80.83 ± 7.88 |
| *Mobility* | **84.17 ± 9.22** | **81.11 ± 8.85** | **82.64 ± 6.55** |
| *Complexity* | 84.17 ± 8.70 | 79.44 ± 10.70 | 81.81 ± 6.67 |

### 3.6 Training with multiple features on active task data – feature fusion (Training Paradigm-II)

We hypothesized in this experiment that different features can provide complementary information and can be combined to enhance the model performance. The top performing features from individual feature training (**Table 2**) – mobility, skew, mean, peak-to-peak (P2P), were used to test the hypothesis. The other three features were not used on the basis that they had a marginally lower specificity. Different approaches to feature fusion were tested in the form of early fusion and late fusion. In the early fusion approach, different features were used as input channels to the best performing network architecture (AT-AR^3^). In late fusion, we tested two different approaches: first, separate time domain models were retrained for each hand-crafted feature, and a single frequency domain module was trained. The domain fusion module was used to combine these time and frequency domain modules (represented as AT-AR^3^-LF^1^). Secondly, we experimented by combining the frequency domain module prior to the feature fusion layer (represented as AT-AR^3^-LF^2^). The performance of these models is presented in **Table 3**.

**Table 3:** Channel classification performance parameters for different approaches using feature fusion from time and frequency domains (training paradigm-II). AT- Active time, AR- autoregressive, EF- early fusion, LF- late fusion.

| Model | Sensitivity %<br>$\mu \pm \sigma$ | Specificity %<br>$\mu \pm \sigma$ | Accuracy %<br>$\mu \pm \sigma$ |
| --- | --- | --- | --- |
| AT-AR<sup>3</sup>-EF | 79.99 ± 9.76 | 83.89 ± 11.37 | 81.94 ± 6.48 |
| AT-AR<sup>3</sup>-LF<sup>1</sup> | 82.22 ± 10.70 | 80.83 ± 9.42 | 81.53 ± 7.43 |
| AT-AR<sup>3</sup>-LF<sup>2</sup> | **85.83 ± 7.80** | **80.27 ± 11.07** | **83.05 ± 6.35** |

### 3.7 Training with individual features on individual stories/active task data (Training Paradigm-III)

In previous experiments so far, we assumed that the channel responds in a similar manner to different stimuli (stories). However, it is plausible that the channel may respond differently to different stimuli. In this training approach, we now assumed that the channel responds differently to different stimuli (stories). The task paradigm consisted of 5 different stories corresponding to 5 different task blocks. We hypothesized that PRCs respond differently to the NRCs for each of these stories. We separated the signals based on the story and train the network in a similar manner as in section 3.6. Each story was trained with its own time domain and frequency domain modules using the AT-AR^3^ network. These different networks were then combined and fed through a fully connected layer. Deep networks with different features as inputs were trained and the performance is compared in **Table 4**, where the features column shows the particular value (mobility, mean, and activity) computed for each story.

**Table 4:** Channel classification performance (mean and variance) for individual active task data, using each story independently (training paradigm-III).

| Features | Sensitivity %<br>$\mu \pm \sigma$ | Specificity %<br>$\mu \pm \sigma$ | Accuracy %<br>$\mu \pm \sigma$ |
| --- | --- | --- | --- |
| *Mobility*<sub>story</sub> | 85.55 ± 10.07 | 79.44 ± 10.91 | 82.5 ± 6.21 |
| *Mean*<sub>story</sub> | 83.61 ± 10.42 | 78.88 ± 11.53 | 81.25 ± 7.35 |
| *Activity*<sub>story</sub> | 83.05 ± 9.73 | 80.55 ± 11.45 | 81.8 ± 6.93 |

## 4. Discussions and Concluding Remarks

In this paper, we proposed novel deep learning architectures to classify the channel response of ECoG signals. The results showed the state-of-the-art classification accuracy of **83.05**% with high specificity and sensitivity of **80.3**% & **85.8**%, respectively, in determining whether the channel was positive (has a response) or negative (has no response) in relation to the task stimulus. The different features and fusion approaches have given us the flexibility in maximizing different metrics, where as an example we can improve the specificity to 83.9% (with a 1% drop in accuracy). In general, with AT-AR^3^-LF^2^ is our best performing model with values >80% for all performance metrics including accuracy, sensitivity and specificity. Traditionally, the accuracy of ECoG-FM is high for mapping sensory and motor function, but relatively low for language modality. On an average, ECoG functional language mapping had a lower sensitivity (62%) and higher specificity (75%) to detect language-specific regions (for a comprehensive review, see Korostenskaja et al. (40)). This is in contrast to the results for hand motor (100% sensitivity and 79.7% specificity) and hand sensory (100% sensitivity and 73.87% specificity) ECoG-based mapping(46). The results of our current study demonstrate that the accuracy for mapping eloquent cortex using ECoG-FM can now be comparable to both sensory and motor ECoG-FM accuracies. The language ECoG-FM accuracy values we have achieved are the highest among those reported so far (47). Although a number of studies have demonstrated successful utilization of the ECoG-FM as a complimentary tool for ESM (13,22). There was not enough evidence to support the use of ECoG-FM as a stand-alone methodology for functional language mapping due to its relatively low accuracy compared with ESM (40,46). The outcome of our research has indicated the potential of ECoG-FM, to be considered as a stand-alone modality for eloquent language cortex localization.

It is possible that some features of the ECoG signal, reflecting the complex nature of language processing, were omitted from consideration when restricting the language ECoG-FM analysis to the gamma frequency band only. Expanding analysis to the whole spectrum of frequencies in our study, therefore, has exceeded the results gained from prior analysis approaches. It contributed to the improved classification accuracy and confirmed the results of previous studies, pointing towards the complex nature of language processing that needs to be considered during the analysis of neurophysiological data. The results of our current study would have a wide-ranging applicability in clinical practice. In particular, our proposed approach can be utilized to prevent functional morbidity post-surgery in patients with pharmacoresistant epilepsy. In addition, this can also be used to increase the accuracy of the eloquent cortex mapping in patients undergoing resection of brain tumours(48) and arteriovenous malformations (49).

Deep learning-based ECoG-FM approaches can also be successfully applied in various fields of adaptive neuro-technologies (e.g., neuromodulation), where ECoG-based mapping is performed to determine the best area for responsive stimulation. For example, for defining the neural correlates of tics, the involuntary movements and/or sounds, to be used for responsive stimulation in patients with Tourette’s syndrome (50,51) and for bidirectional neurostimulation via fully implantable neural interfaces in Parkinson’s disease (52). The applicability of our proposed approach extends as well towards the fields of developmental disorders (e.g., autism) (53,54), psychiatry (e.g., major depression (55) and obsessive-compulsive disorder (OCD) (56), addiction (57), eating disorders and obesity (58,59), where neurostimulation can be potentially utilized to provide treatment and improve patients’ quality of life.

### 4.1 Why Deep Learning in Studies with Limited Subjects

It is traditionally argued that deep learning models are data intensive and only fit to problems where adequate training data is available. The inherently low dimensional nature of physiological data (such as ECoG and EEG) both in terms of samples and subjects has historically restricted this field from taking advantage of the recent advances in machine learning (led by novel deep learning architectures). This trend is arguable for multiple reasons. The number of subjects (which in most of these studies is low) is not a good parameter to decide whether we can use deep learning-based methods. Our results clearly support this argument, where the number of subjects (11 for training the models) could seem small. But the overall data (from 128 electrodes at 1200 samples per second) having millions of samples was highly sufficient to efficiently train our deep learning models. Our classification accuracy (83.05%) and sensitivity (85.8%) bodes well for selecting deep learning methods in our proposed models. This trend can also be seen in other recent studies^52-55^ and we are observing a paradigm shift in classification tasks using 1D physiological data. At the same time, deep learning models are developed that can work with small data. In particular, Bayesian deep learning models are shown to have good performance even when the labelled training data is scarce^55^. We conclude that even with limited subjects, the data could be sufficient for successfully using deep learning methods in 1D data classification tasks such as ECoG-FM.

### 4.2 Limitations and future perspectives

Despite the state-of-the-art results in ECoG-FM predictions, there are some limitations of our work to be noted. First, our experimental paradigm (**Figure 4**) involves five different stories being played to the subject. The responses to these stories, have some inherent similarities, but overall are different. Therefore, training the deep learning model with a single label for the whole channel could add noise to the model. Though we have tested the effect of training the network, while treating each story individually, this reduces the overall data available to train the model. We believe that these results can be improved by including additional data and then training the system individually for each story in the paradigm. Second, the subjects used in this initial validation study were a mix of teenagers and adults. Smith^56^ found that the effect of epilepsy and seizures on children and adults was different i.e., the rules learned about the behavior of the brain in adults is different for children. Hence, a more comprehensive study with focus on children/teenagers is needed. This is one of our future aims to test the proposed machine learning based approach in different patient populations; however, patient recruitment is difficult due to the involvement of surgery, and disease prevalence.

Finally, in the proposed approach, due to the exploratory and research nature of our study, the classification was not performed in real-time. We have used retrospective data for validating the innovations and are currently working on the real-time clinical implementation of the algorithm. We intend to extend this study for mapping functional language cortex in prospective subjects. We believe that implementing such a reliable technology will increase current presurgical and intra-operative functional mapping accuracy, expand surgical treatment opportunities, prevent post-surgical language morbidity, and improve patient outcomes.

## Acknowledgements

The authors acknowledge The Central Florida Health Research (CFHR) grant for supporting this study.

## Authors’ Contributions

Idea development, current study conception: MK, UB, EMC.

Deep learning design, computational experiments: HR, CS, SMA, UB.

Patient recruitment, IRB process, ECoG tasks, ESM experiments: MK, CMS, EMC, and KHL.

Manuscript writing, editing, and significant revision: All authors.

## References

1. Patricia S. About Epilepsy: The Basics [Internet]. Epilepsy Foundation; 2018. Available from: https://www.epilepsy.com/learn/about-epilepsy-basics

2. Penfield W, Rasmussen T. The cerebral cortex of man; a clinical study of localization of function. 1950;

3. Penfield W, Roberts L. Mapping the speech area. Speech brain Mech. 1959;103–18.

4. Ojemann G, Ojemann J, Lettich E, Berger M. Cortical language localization in left, dominant hemisphere: an electrical stimulation mapping investigation in 117 patients. J Neurosurg. 1989;71(3):316–26.

5. Ojemann JG, Ojemann GA, Lettich E. Cortical stimulation mapping of language cortex by using a verb generation task: effects of learning and comparison to mapping based on object naming. J Neurosurg. 2002;97(1):33–8.

6. Edwards E, Nagarajan SS, Dalal SS, Canolty RT, Kirsch HE, Barbaro NM, et al. Spatiotemporal imaging of cortical activation during verb generation and picture naming. Neuroimage. 2010;50(1):291–301.

7. Pouratian N, Cannestra AF, Bookheimer SY, Martin NA, Toga AW. Variability of intraoperative electrocortical stimulation mapping parameters across and within individuals. Journal of Neurosurgery Publishing Group; 2004.

8. Bank AM, Schevon CA, Hamberger MJ. Characteristics and clinical impact of stimulation-evoked seizures during extraoperative cortical mapping. Epilepsy Behav. 2014;34:6–8.

9. Schalk G, Leuthardt EC, Brunner P, Ojemann JG, Gerhardt LA, Wolpaw JR. Real-time detection of event-related brain activity. Neuroimage. 2008;43(2):245–9.

10. Leuthardt EC, Miller K, Anderson NR, Schalk G, Dowling J, Miller J, et al. Electrocorticographic frequency alteration mapping: a clinical technique for mapping the motor cortex. Oper Neurosurg. 2007;60(suppl_4):ONS--260.

11. Miller KJ, Shenoy P, Miller JW, Rao RPN, Ojemann JG, others. Real-time functional brain mapping using electrocorticography. Neuroimage. 2007;37(2):504–7.

12. Kapeller C, Korostenskaja M, Prueckl R, Chen P-C, Lee KH, Westerveld M, et al. CortiQ-based real-time functional mapping for epilepsy surgery. J Clin Neurophysiol. 2015;32(3):e12.-e22.

13. Prueckl R, Kapeller C, Gruenwald J, Ogawa H, Kamada K, Korostenskaja M, et al. Passive functional mapping guides electrical cortical stimulation for efficient determination of eloquent cortex in epilepsy patients. In: 2017 39th Annual International Conference of the IEEE Engineering in Medicine and Biology Society (EMBC). 2017. p. 4163–6.

14. Towle VL, Syed I, Berger C, Grzesczcuk R, Milton J, Erickson RK, et al. Identification of the sensory/motor area and pathologic regions using ECoG coherence. Electroencephalogr Clin Neurophysiol. 1998;106(1):30–9.

15. Qian T, Zhou W, Ling Z, Gao S, Liu H, Hong B. Fast presurgical functional mapping using task-related intracranial high gamma activity. J Neurosurg. 2013;119(1):26–36.

16. Vansteensel MJ, Bleichner MG, Dintzner LT, Aarnoutse EJ, Leijten FSS, Hermes D, et al. Task-free electrocorticography frequency mapping of the motor cortex. Clin Neurophysiol. 2013;124(6):1169–74.

17. Alkawadri R, Gaspard N, Hirsch LJ, Spencer DD. T105. A novel method for ECOG-based localization of function. Clin Neurophysiol. 2018;129:e43.

18. Prueckl R, Kapeller C, Potes C, Korostenskaja M, Schalk G, Lee KH, et al. cortiQ-Clinical software for electrocorticographic real-time functional mapping of the eloquent cortex. In: Engineering in Medicine and Biology Society (EMBC), 2013 35th Annual International Conference of the IEEE. 2013. p. 6365–8.

19. Arya R, Wilson JA, Vannest J, Byars AW, Greiner HM, Buroker J, et al. Electrocorticographic language mapping in children by high-gamma synchronization during spontaneous conversation: Comparison with conventional electrical cortical stimulation. Epilepsy Res [Internet]. 2015;110:78–87. Available from: http://www.sciencedirect.com/science/article/pii/S0920121114003271

20. Arya R, Wilson JA, Fujiwara H, Vannest J, Byars AW, Rozhkov L, et al. Electrocorticographic high-gamma modulation with passive listening paradigm for pediatric extraoperative language mapping. Epilepsia. 2018;59(4):792–801.

21. Korostenskaja M, Kamada K, Guger C, Salinas CM, Westerveld M, Castillo EM, et al. Electrocorticography-Based Real-Time Functional Mapping for Pediatric Epilepsy Surgery. J Pediatr Epilepsy. 2015;4(04):184–206.

22. Prueckl R, Kapeller C, Gruenwald J, Guger C, Fernandes F, Walchshofer M, et al. O202 Combining the strengths of passive functional mapping and electrical cortical stimulation. Clin Neurophysiol. 2017;128(9):e243.

23. RaviPrakash H, Korostenskaja M, Lee K, Baumgartner J, Castillo E, Bagci U. Automatic response assessment in regions of language cortex in epilepsy patients using ECoG-based functional mapping and machine learning. In: Systems, Man, and Cybernetics (SMC), 2017 IEEE International Conference on. 2017. p. 519–24.

24. Korostenskaja M, RaviPrakash H, Bagci U, Lee KH, Chen PC, Salinas C, et al. Gold Standard for epilepsy/tumor surgery coupled with deep learning offers independence to a promising functional mapping modality. In: Guger C, Allison B, Ushiba J, editors. Brain-Computer Interface Research: A State-of-the-Art Summary 6. Springer International Publishing: Cham; 2017.

25. Graves A, Schmidhuber J. Framewise phoneme classification with bidirectional LSTM and other neural network architectures. Neural Networks. 2005;18(5–6):602–10.

26. Baccouche M, Mamalet F, Wolf C, Garcia C, Baskurt A. Spatio-Temporal Convolutional Sparse Auto-Encoder for Sequence Classification. In: BMVC. 2012. p. 1–12.

27. Wang J, Yang Y, Mao J, Huang Z, Huang C, Xu W. Cnn-rnn: A unified framework for multi-label image classification. In: Proceedings of the IEEE conference on computer vision and pattern recognition. 2016. p. 2285–94.

28. Inuso G, La Foresta F, Mammone N, Morabito FC. Brain activity investigation by EEG processing: Wavelet analysis, kurtosis and Renyi’s entropy for artifact detection. Proc 2007 Int Conf Inf Acquis ICIA. 2007;195–200.

29. Pollock VE, Schneider LS, Lyness SA. EEG amplitudes in healthy, late-middle-aged and elderly adults: normality of the distributions and correlations with age. Electroencephalogr Clin Neurophysiol. 1990;75(4):276–88.

30. Hjorth B. EEG analysis based contributions on time domain properties. Electroencephalogr Clin Neurophysiol [Internet]. 1970;306–10. Available from: http://ac.els-cdn.com.ezlibproxy1.ntu.edu.sg/0013469470901434/1-s2.0-0013469470901434-main.pdf?_tid=44c4f902-55ad-11e7-ae24-00000aacb35f&acdnat=1497959013_c82140ec01fb64b12901b87d9aba3f05

31. Goodfellow SD, Goodwin A, Greer R, Laussen PC, Eytan D, Goodfellow SD, et al. Towards Understanding ECG Rhythm Classification Using Convolutional Neural Networks and Attention Mappings ECG Classification Using Convolutional Neural Networks and Attention Mappings. Mlhc-2018 [Internet]. 2018;85(August):1–18. Available from: https://static1.squarespace.com/static/59d5ac1780bd5ef9c396eda6/t/5b73729d562fa79aabf87383/1534292642748/9.pdf

32. Zabihi M, Rad AB, Kiranyaz S, Särkkä S, Gabbouj M. 1D Convolutional Neural Network Models for Sleep Arousal Detection. 2019;(Schulz 2008):1–10. Available from: http://arxiv.org/abs/1903.01552

33. Hochreiter S, Schmidhuber J. Long short-term memory. Neural Comput. 1997;9(8):1735–80.

34. Liu Q, Zhou F, Hang R, Yuan X. Bidirectional-Convolutional LSTM Based Spectral-Spatial Feature Learning for Hyperspectral Image Classification. arXiv Prepr 170307910. 2017;

35. Nowak J, Taspinar A, Scherer R. LSTM Recurrent Neural Networks for Short Text and Sentiment Classification. In: International Conference on Artificial Intelligence and Soft Computing. 2017. p. 553–62.

36. Flunkert V, Salinas D, Gasthaus J. DeepAR: Probabilistic Forecasting with Autoregressive Recurrent Networks. arXiv Prepr 170404110. 2017;

37. Gammulle H, Denman S, Sridharan S, Fookes C. Two Stream LSTM: A Deep Fusion Framework for Human Action Recognition. In: Applications of Computer Vision (WACV), 2017 IEEE Winter Conference on. 2017. p. 177–86.

38. Lin M, Chen Q, Yan S. Network in network. arXiv Prepr 13124400. 2013;

39. Korostenskaja M, Chen P-C, Salinas CM, Westerveld M, Brunner P, Schalk G, et al. Real-time functional mapping: potential tool for improving language outcome in pediatric epilepsy surgery: Case report. J Neurosurg Pediatr. 2014;14(3):287–95.

40. Korostenskaja M, Wilson AJ, Rose DF, Brunner P, Schalk G, Leach J, et al. Real-time functional mapping with electrocorticography in pediatric epilepsy: comparison with fMRI and ESM findings. Clin EEG Neurosci. 2014;45(3):205–11.

41. Karim F, Majumdar S, Darabi H, Chen S. LSTM Fully Convolutional Networks for Time Series Classification. IEEE Access. 2017;6:1662–9.

42. Karim F, Majumdar S, Darabi H, Harford S. Multivariate LSTM-FCNs for time series classification. Neural Networks. 2019;116:237–45.

43. Dietterich T. Overfitting and undercomputing in machine learning. ACM Comput Surv. 1995;27(3):326–7.

44. James G, Witten D, Hastie T, Tibshirani R. An introduction to statistical learning. Vol. 112. Springer; 2013.

45. Arlot S, Celisse A. A survey of cross-validation procedures for model selection. Stat Surv. 2010;4:40–79.

46. Kapeller C, Korostenskaja M, Prueckl R, Chen P-C, Lee KH, Westerveld M, et al. CortiQ-based Real-Time Functional Mapping for Epilepsy Surgery. J Clin Neurophysiol. 2015;32(3):e12.-e22.

47. Arya R, Horn PS, Crone NE. ECoG high-gamma modulation versus electrical stimulation for presurgical language mapping. Epilepsy Behav. 2018;79:26–33.

48. Ogawa H, Kamada K, Kapeller C, Hiroshima S, Prueckl R, Guger C. Rapid and minimum invasive functional brain mapping by real-time visualization of high gamma activity during awake craniotomy. World Neurosurg. 2014;82(5):912--e1.

49. Kamada K, Ogawa H, Saito M, Tamura Y, Anei R, Kapeller C, et al. Novel techniques of real-time blood flow and functional mapping. Neurol Med Chir (Tokyo). 2014;54(10):775–85.

50. Molina R, Okun MS, Shute JB, Opri E, Rossi PJ, Martinez-Ramirez D, et al. Report of a patient undergoing chronic responsive deep brain stimulation for Tourette syndrome: proof of concept. J Neurosurg. 2018;129(2):308–14.

51. Shute JB, Okun MS, Opri E, Molina R, Rossi PJ, Martinez-Ramirez D, et al. Thalamocortical network activity enables chronic tic detection in humans with Tourette syndrome. NeuroImage Clin. 2016;12:165–72.

52. Swann NC, de Hemptinne C, Miocinovic S, Qasim S, Ostrem JL, Galifianakis NB, et al. Chronic multisite brain recordings from a totally implantable bidirectional neural interface: experience in 5 patients with Parkinson’s disease. J Neurosurg. 2018;128(2):605–16.

53. Sinha S, McGovern RA, Sheth SA. Deep brain stimulation for severe autism: from pathophysiology to procedure. Neurosurg Focus. 2015;38(6):E3.

54. Sturm V, Fricke O, Bührle CP, Lenartz D, Maarouf M, Treuer H, et al. DBS in the basolateral amygdala improves symptoms of autism and related self-injurious behavior: a case report and hypothesis on the pathogenesis of the disorder. Front Hum Neurosci. 2013;6:341.

55. Coenen VA, Sajonz B, Reisert M, Bostroem J, Bewernick B, Urbach H, et al. Tractography-assisted deep brain stimulation of the superolateral branch of the medial forebrain bundle (slMFB DBS) in major depression. NeuroImage Clin. 2018;20:580–93.

56. Vázquez-Bourgon J, Martino J, Sierra MP, Infante JC, Mart\’\inez MÁM, Ocón R, et al. Deep brain stimulation and treatment-resistant obsessive-compulsive disorder: A systematic review. Rev Psiquiatr Salud Ment. 2017;

57. Wang TR, Moosa S, Dallapiazza RF, Elias WJ, Lynch WJ. Deep brain stimulation for the treatment of drug addiction. Neurosurg Focus. 2018;45(2):E11.

58. Val-Laillet D, Aarts E, Weber B, Ferrari M, Quaresima V, Stoeckel LE, et al. Neuroimaging and neuromodulation approaches to study eating behavior and prevent and treat eating disorders and obesity. NeuroImage Clin. 2015;8:1–31.

59. Oterdoom DLM, van Dijk G, Verhagen MHP, Jiawan VCR, Drost G, Emous M, et al. Therapeutic potential of deep brain stimulation of the nucleus accumbens in morbid obesity. Neurosurg Focus. 2018;45(2):E10.

