## Supplementary Information for "Deep Learning provides exceptional accuracy to ECoG-based Functional Language Mapping for epilepsy surgery"

### Supplementary Information (SI)

#### *Patient Demographics*

**Supplementary Table 1:** Patient demographics, clinical information, grid placement, and information about the number of analysed channels (electrodes) are summarized. All study participants were left hemisphere language dominant for language.

| Subject # | Age (yrs) | Sex* | Epilepsy Focus | Grid Placement | Epilepsy Onset (yrs) | All Tested Channels/ PRC / NRC** |
|---|---|---|---|---|---|---|
| 1 | 19 | M | Frontal-Temporal | Left | 16 | 54 / 22 / 32 |
| 2 | 33 | F | Frontal-Temporal | Left | 10 | 32 / 5 / 27 |
| 3 | 20 | M | Frontal-Temporal | Left | 6 | 127 / 16 / 111 |
| 4 | 22 | F | Parietal | Left | 20 | 30 / 19 / 11 |
| 5 | 32 | F | Temporal | Bilateral | 26 | 48 / 10 / 38 |
| 6 | 52 | M | Temporal | Left | 30 | 48 / 5 / 43 |
| 7 | 15 | M | Temporal-Parietal | Left | 12 | 43 / 10 / 33 |
| 8 | 17 | M | Frontal-Temporal | Left | 7 | 72 / 20 / 52 |
| 9 | 15 | F | Medial-Frontal | Left | 7 | 80 / 19 / 61 |
| 10 | 17 | M | Frontal-Temporal | Left | 16 | 63 / 10 / 53 |
| 11 | 13 | F | Temporal | Left | 2 | 40 / 6/ 34 |

**F – Female, M - Male*

***PRC - positive response channels, NRC - negative response channels.*

#### *Pre-processing ECoG Signal*

We recorded the signals from the ECoG grid electrodes for a few seconds prior to the task initiation. Similarly, we recorded the signals for a few seconds post task completion. This led to padding of data, that may not contain useful information but was necessary to ensure that signals were located at the boundary locations. Thus, as a first step of preparing the recorded data for machine learning (ML) based analysis, non-task/control time points in the signal were eliminated: these correspond to the spontaneous activity recording before the 0-min and any trailing signals at the end of the experiment. We recorded the ECoG signals over a few minutes at a sampling rate of $\sim$ 1200Hz, that was a few hundred thousand data points per electrode (1200 *samples/s* $*$ 300*s*). Due to this large sample size, we used the sliding window approach to generate features and analyzed the whole signal through these feature representations. One drawback of this approach might be a potential loss of long term dependencies; however, we believe that we have minimized the potential loss of periodic information because of the paradigm involving different stories and a wise choice of data preparation.

##### *Estimating autoregressive (AR) parameters*

Methods to solve for the AR parameters are diverse and can be classified into three main categories(1):

1. *Correlation Function Estimation method* - AR parameter estimation from autocorrelation sequence estimates of Yule-Walker equations(1). These equations relate the autocorrelation sequence for lags $0$ to $p$, to the variance of a driving noise signal. This approach has the benefit of producing a stable model, however, in some special cases, involving nearly periodic signals, it may produce incorrect estimates.

2. *Reflection coefficient estimation methods* - AR parameters are constrained to satisfy a recursive relationship. The Levinson-Durbin recursion(1) relates the AR parameters of order $p$ to the AR parameters of order $p - 1$ as
$$a_p[n] = a_{p-1}[n] + k_p a^*_{p-1}[p-n] \qquad (1)$$
where, $k_p$ is the reflection coefficient.
The benefit of this approach is that it produces stable models.

3. *Least Squares Linear Prediction Estimation methods* - AR parameters are estimated by the minimization of the forward (predicting the future) and backward (predicting the past) linear prediction squared errors. The minimization can be done separately or combined. The advantage of this approach is that it minimizes the residual variance however, this does not directly translate to minimizing the variance of the prediction error.

We used the reflection coefficient estimation-based methods, which require the estimation of $k_p$ from the autocorrelation function for lags $0$ to $p-1$. These approaches are based on minimizing the least square error of the forward (predicting the future) and backward (predicting the past) linear prediction with respect to the reflection coefficient. Other known approaches in this category includes the geometric algorithm, which aims to minimize the geometric mean of the squared error predictions, the harmonic algorithm that minimizes the arithmetic mean instead of the geometric mean, and the maximum likelihood approach. Among these solutions, we used the popular Harmonic algorithm, also known as Burg method(1) which produces comparable estimates to the least square linear prediction estimation method. In the Burg estimation approach, the reflection coefficient $k_p$ was found by,

$$k_p = \frac{-2\sum_{n=p+1}^{N} e_{p-1}^{f}[n]\, e_{p-1}^{b*}[n-1]}{\sum_{n=p+1}^{N}\left|e_{p-1}^{f}[n]\right|^2 + \sum_{n=p+1}^{N}\left|e_{p-1}^{b}[n-1]\right|^2} \quad (2)$$

where $e_p^f, e_p^b$ are the forward and backward prediction errors and are expressed as

$$e_p^f[n] = e_{p-1}^f[n] + k_p e_{p-1}^b[n-1] \quad (3)$$

$$e_p^b[n] = e_{p-1}^b[n-1] + k_p^* e_{p-1}^f[n] \quad (4)$$

Once the errors in Eqs. (3)~~(3)~~ and (4)~~(4)~~ were computed recursively, the reflection coefficient was computed by minimizing the arithmetic mean of the squared forward and backward prediction errors. This allowed us to solve for the final AR parameters, $a_p[k]$ in Eq. (1)~~(1)~~, which were used to characterize the ECoG signals from frequency-domain perspective.

***Time Domain Features***

The following time domain features were used as input to our deep learning architectures.

1. Mean – The mean signal intensity within the window of a channel's signal block is used as a feature. This can help detect the change in activation across windows and is measured as:

   $$m_i = \frac{\sum_{j=t}^{t+n} X_j}{n},$$

   where *i*=*i*th window, *n*=size of window, *t*=time point in the window of channel signal *X*
2. Skew – This is a measure of the symmetry in a distribution and is measured as

   $$s_i = \frac{\sum_{j=t}^{t+n}\left(X_j - m_i\right)^3}{\sigma^3},$$

   where $\sigma$ is the standard deviation within the window *i*.
3. Kurtosis – This is a measure of how peaked around the mean a distribution is and is measured as

   $$k_i = \frac{\sum_{j=t}^{t+n}\left(X_j - m_i\right)^4}{\sigma^4}$$
4. Peak-to-peak (P2P) – This measures the difference between the minimum and maximum values in the distribution and is measured as

   $$p2p_i = \max X_{t:t+n} - \min X_{t:t+n}$$
5. Hjorth features – define the signal in terms of amplitude, time scale and complexity.
   a. Activity – this is a measure of the mean power of the signal and is computed as the variance of the signal.
   b. Mobility – this represents the mean frequency and is defined as the square root of the ratio of the variance of the first derivative and the amplitude.

c. Complexity – this represents the frequency change and is defined as the ratio between the mobility of the first derivative of the signal and the mobility of the signal itself.

***Data Preparation***

We generated a new time-series data from the existing one as follows: for a time-series ECoG signal $T = \{t_1, t_2, ..., t_n\}$, starting at a particular time $t$, with a window size $k$ and stride $s$, $\frac{n-k}{s} + 1$, sub-blocks of time-series signals were generated by sliding the window across the channel's recorded signal. In our implementation, step-size $s$, is chosen as 100 to generate fewer redundant samples.

From **Figure 2** (main text), we have 10 blocks of ECoG data (active task + control) for each electrode. This amounts to a recording of 360,000 samples per channel ($10 * 30s * 1200 samples/s = 360{,}000$). Following basic data pre-processing steps, a sliding window of width 600 samples (i.e. 0.5 sec) and a stride of 100 samples are used on each data block (active/control). This yields a total of 354 sub-blocks ($\frac{n-k}{s} + 1 = \frac{30s*1200samples/s-600samples}{100samples} = 354$). Features are extracted from each of these sub-blocks and concatenated to generate a single feature vector per channel. The proposed system is trained on these blocks of data and the final ECoG signal classification involves a basic majority voting approach over these (Step 5).

In this study, our focus was not to explore different data augmentation techniques, but to simply maximize the data size to fit the model. Instead of the method that we used for data augmentation, different techniques may be used, such as the method by Le Guennec et al.(2) where randomly selected slices were warped to speed up or slow down the signal followed by the window slicing approach and by DeVries and Taylor(3), where simple transformations were applied to existing data points. More advanced data augmentation approaches in the form of transfer learning(4–6) and generative adversarial networks(7) may also be considered.

***Computational Environment and Network Parameters***

Prediction models were trained and tested using Keras with TensorFlow backend on servers equipped with NVidia Titan X with 12Gb Graphics memory, 2.7GHz CPU, 64 GB RAM. Data were stored as '.mat' files after reading the data using EEGLAB(8). Python 2.7 was used to analyze the signals in a fully automated fashion. We have used the following network architecture design parameters for optimal prediction of the channel response (**Supplementary Table 2**). As common to neural network architecture design, these architecture design parameters were found empirically and with thorough comparisons with alternative parameter selections.

**Supplementary Table 2:** Network design parameters - $AT\text{-}AR^4$.

| Layer | Filter Size |
|---|---|
| *1D Convolution (Time Domain RNN)* | 128, 64, 128 |
| *Dense (Frequency Domain Features)* | 64 |
| *LSTM (Time Domain RNN)* | 8 |
| *Dense (Domain Fusion Network)* | 64, 32, 2 |
| Global Hyperparameters | Value |
| *Loss* | Categorical Cross-Entropy |
| *Optimizer* | Adam |
| *Learning Rate* | 0.001 |
| *Exponential decay rate 1* | 0.9 |
| *Exponential decay rate 2* | 0.999 |
| *Decay every 'n' epochs* | 25 |

#### References for Supplementary Information


1. Marple SL. Digital spectral analysis: with applications. Vol. 5. Prentice-Hall Englewood Cliffs, NJ; 1987.

2. Le Guennec A, Malinowski S, Tavenard R. Data Augmentation for Time Series Classification using Convolutional Neural Networks. In: ECML/PKDD Workshop on Advanced Analytics and Learning on Temporal Data. 2016.

3. DeVries T, Taylor GW. Dataset augmentation in feature space. arXiv Prepr arXiv170205538. 2017;

4. LeCun Y, Bengio Y, Hinton G. Deep learning. Nature. 2015;521(7553):436.

5. Jayaram V, Alamgir M, Altun Y, Scholkopf B, Grosse-Wentrup M. Transfer learning in brain-computer interfaces. IEEE Comput Intell Mag. 2016;11(1):20–31.

6. Perez L, Wang J. The effectiveness of data augmentation in image classification using deep learning. arXiv Prepr arXiv171204621. 2017;

7. Esteban C, Hyland SL, Rätsch G. Real-valued (medical) time series generation with recurrent conditional GANs. arXiv Prepr arXiv170602633. 2017;

8. Delorme, Arnaud and Makeig S. EEGLAB: an open source toolbox for analysis of single-trial EEG dynamics including independent component analysis. J Neurosci Methods. 2004;134(1):9--21.
